# Disrupting pathogenic interactions between α-synuclein, c-Abl, and redox stress

**DOI:** 10.1101/840306

**Authors:** Soumitra Ghosh, Seok Joon Won, Rebecca Fong, Nicholas J. M. Butler, Arianna Moss, Candance Wong, June Pan, Jennifer Sanchez, Long Wu, Jiejie Wang, Fredric P. Manfredsson, Raymond A. Swanson

## Abstract

**Objective:** Redox stress, c-Abl activation, and α-synuclein aggregates each independently contribute to neurodegeneration in Parkinson’s disease. Interactions between these factors may underlie convergent and feed-forward mechanisms of disease progression.

**Methods:** α-synuclein aggregate formation was induced in neuronal cultures and mouse substantia nigra by exposure to pre-formed human α-synuclein fibrils or by AAV-mediated over-expression of α-synuclein. Aggregate formation, c-Abl activation, redox stress, and neurodegeneration were evaluated by immunohistochemistry and Western blots, and mouse motor function was evaluated using the rota-rod and pole tests. To suppress redox stress, cultures and mice were treated with N-acetyl cysteine, a thiol repletion agent that supports neuronal glutathione metabolism.

**Results:** In primary neuron cultures, the formation of α-synuclein aggregates led to redox stress and c-Abl activation. Redox stress alone, in the absence of α-synuclein aggregates, was also sufficient to induced c-Abl activation. N-acetyl cysteine suppressed redox stress, and likewise suppressed both c-Abl activation and α-synuclein aggregation. A similar pattern was observed in the two mouse models of Parkinson’s disease. In both models, α-synuclein aggregates in the substantia nigra were accompanied by redox stress, c-Abl activation, dopaminergic neurodegeneration and motor impairment, all of which were attenuated in mice treated with oral N-acetyl cysteine.

**Interpretation:** These results indicate that α-synuclein aggregates induce c-Abl activation by a redox stress mechanism. c-Abl in turn promotes α-synuclein aggregation, and this potentially feed-forward process can be blocked by N-acetyl cysteine. The findings thus add mechanistic support for N-acetyl cysteine as a therapeutic for Parkinson’s disease.

## INTRODUCTION

Risk factors for Parkinson’s disease (PD) include exposure to toxins and environmental agents that affect cell redox state, such as MPTP, rotenone, paraquat, and others ^1-3^. Risk for PD is also increased by genetic variants that promote formation of α-synuclein aggregates ^4^. These environmental and genetic factors can interact, as redox state influences α-synuclein aggregation ^3, 5, 6^ and, conversely, α-synuclein aggregates induce oxidative stress ^7, 8^. These bidirectional interactions suggest the possibility of convergent or feed-forward relationships that could be targeted to suppress disease progression. However, the mechanisms by which these factors interact *in vivo* is not well understood.

The tyrosine kinase c-Abl provides one potential link between cell redox state and α-synuclein aggregation. c-Abl is a ubiquitous non-receptor protein kinase. It normally functions as part of the cellular DNA damage response ^9^, but mutations in c-Abl that cause constitutive activation contribute to myeloproliferative and other cancer types ^10^. Cell culture studies suggest that c-Abl can be activated by oxidative stress, although it is uncertain if this is a direct effect or an indirect effect of DNA damage ^11-13^. In animal models of PD, c-Abl activation promotes both α-synuclein aggregate formation and neuronal death ^13, 14^. The effect of c-Abl on aggregate formation is mediated in part by its phosphorylation of the ubiquitin ligase, parkin, as phosphorylated parkin is unable to ubiquitinate substrates such as α-synuclein and thus slows their normal proteolysis ^15, 16^. c-Abl may also promote aggregate formation through direct phosphorylation of α-synuclein at residue 39 ^14, 17^. Pharmacological inhibition of c-Abl can slow disease progression in animal models of PD ^13, 18^, and a clinical trial of c-Abl inhibitor, nilotinib, is currently underway. However, nilotinib and other c-Abl inhibitors as a class exhibit significant cardiac, gastrointestinal and other toxicities ^19^, which may limit their use in chronic diseases such as PD.

Here we sought to determine whether c-Abl activation mediates interaction between redox stress and α-synuclein aggregate formation, and whether suppressing redox stress can suppress the effects of c-Abl on α-synuclein aggregation. We used cell culture and two mouse models of PD; stereotactic injection of pre-formed α-synuclein fibrils, and AAV-mediated α-synuclein overexpression. Our findings show that α-synuclein aggregates activate c-Abl through an oxidant stress mechanism that can be blocked by the thiol agent N-acetyl cysteine (NAC). Surprisingly, NAC also suppressed formation of α-synuclein aggregates. The results identify a feed-forward mechanism by which α-synuclein aggregates can drive c-Abl activation, and support accumulating evidence that N-acetyl cysteine could be an effective therapeutic intervention for PD.

## MATERIALS and METHODS

All studies were performed in accordance with the US PHS Policy on Humane Care and Use of Laboratory Animals and with protocols approved by the San Francisco Veterans Affairs Medical Center animal studies committee. Cell culture reagents were obtained from Sigma-Aldrich except where noted. The primary antibodies and concentrations used for western blotting (WB) and immunocytochemistry (IHC) are as follows: c-Abl, AbCAM-ab15130 (WB 1:2000); phospho c-Abl 245, Cell Signaling-2861 (WB 1:1000, IHC 1:250); α-synuclein, LifeTech-328100 (WB 1:1000); phospho synuclein 39, BioLegend A15119B (WB 1:500, IHC 1:200); phospho synuclein 129, AbCAM-ab59264 (IHC 1:1000) and AbCAM-ab184674 (IHC 1:500); 4-hydroxynonenal, Alpha Diagnostics 13-M (IHC 1:1000) and 12-S (IHC 1:2500); microtubule associated protein 2, Millipore MAB3418 (IHC 1:2000); tyrosine hydroxylase, Millipore AB1542 (IHC 1:2000); β-actin, ThermoFisher MA1-140 (WB 1:5000).

### Pre-formed α-synuclein fibrils (PFFs)

Pre-formed human α-synuclein fibrils were prepared from recombinant human α-synuclein as described ^20^ and stored at -80 °C. Aliquots were thawed and sonicated for 5 minutes immediately before use.

### rAAV production

Recombinant AAV genomes were generated by inserting full length human synuclein or mCherry behind the CAG (CBA/CMV hybrid) promoter and packaged in rAAV2/9 as described ^21^. In brief, 293T cells were transfected with genome and helper plasmids. Three days later, virus was harvested from media and cells and purified using an idodixanol gradient. Virus was quantified using digital droplet PCR, and titers were diluted to 1.0 × 10^13^ vector genomes / ml.

### Cell culture studies

Neurons were isolated from embryonic day 17 / 18 of C57BL/6 wildtype or α, β,γ synuclein triple knockout mice ^22^ and plated onto poly-D-lysine coated glass coverslips as described ^23^. The cultures were maintained in medium containing 5 mM glucose, with Neurobasal (Life technologies #21103049), and B27 (Life technologies #17504044) supplements, in a 5% CO_2_, 37 °C incubator. Exposure to PFFs was performed by adding 1 µg / ml to culture media of neurons on day 7 *in vitro*. Where used, N-acetyl cysteine (NAC, 400 µM) was added on day 9 *in vitro* at a 400 µM concentration. Immunostaining and live-cell assessment of reactive oxygen species with dihydroethidium were performed as described ^23^. Three to five photos were taken of each coverslip by an observer blinded to the experimental conditions, and with identical camera settings maintained within each experiment. Fluorescence intensity was measured as arbitrary units and normalized to either the number of cells or neuronal area in each field. For live-cell assessment of reactive oxygen species, 10 µM dihydroethidium (Life Technologies, D11347) was added to the culture medium 30 minutes prior to fixation.

Cells were lysed in Phospho-Sure lysis buffer and the tissue lysate was then sonicated on ice, agitated on a rotator at 4 °C for 1 hour, and centrifuged for 30 minutes at 4°C, 10,000 *g.* Proteins in the supernatant were separated by electrophoresis and transferred to PVDF membranes. The membranes were then blocked with 5% non-fat milk or goat serum, incubated with primary antibody overnight, with secondary antibody (IR 600/800 CW, Licor, #C70908-04 or #C70620-05) for 1 hour, and imaged on an Odyssey gel scanner. Where chemiluminescence imaging was used, membranes were incubated with secondary antibodies (Sigma, Rabbit-GENA934 and Mouse-GENA931) and treated with Pierce Super signal reagent (ThermoFisher #34580). For immunoblots of α-synuclein oligomers ^24^,1% SDS was added to the cell lysates, the lysates were centrifuged at 16,000 *g*, and the PVDF membranes were treated with paraformaldehyde prior to immunoblotting ^24^.

### Mouse studies

The *in vivo* studies used either wild-type C57Bl/6 mice or EAAC1^-/-^ male mice ^25^ on the C57BL/6 background. Mice were arbitrarily assigned to treatment groups at age 9 months and euthanized at age 15 months. Where used, N-acetyl cysteine (NAC) was provided in drinking water at a concentration of 3 mg / ml, and exchanged twice weekly with fresh solution. Of the 26 mice used for the studies there were two premature mortalities, both in the AAV-synuclein group.

### Stereotactic injections

For AAV injections, a microinjection syringe was inserted to target the substantia nigra unilaterally (anterior–posterior, +/-3.0, medio-lateral, +1.5, dorso-ventral, -4.6 from bregma). 5 μl of AAV2/9 encoding either human α-synuclein or mCherry was injected at a concentration of 1.0 × 10^13^ vector genomes / ml), at a flowrate of 0.25 µl / minute ^26^. Post-surgical incisional pain was treated with bupivacaine and buprenorphine. Stereotactic PFF injections were performed in the same way but were bilateral. Each injection delivered 10 µl of 5 μg / μl PFF or, for controls, 10 µl of PBS.

### Behavioral assessments

The rota-rod test and pole tests were performed as described ^27, 28^, with minor modifications. Two persons handled the mice, and both were blinded to the treatment group assignments. For the rota-rod test, mice were first habituated to the task by placement for 90 seconds on a horizontal rod (San Diego Instruments Rota-rod) rotating at 4 rpm (1^st^ day) and 5.5 rpm (2^nd^ day), 5 times each day. For testing, mice were placed on the horizontal rod at a start speed 0 rpm and an acceleration rate of 6 rpm / minute, and the time until fall from the rod was recorded. Tests were repeated 6 times each day for 2 consecutive days. The outlier (low value) for each mouse was discarded and the average time from the remaining 11 trials was recorded for each mouse. For the pole test, mice were placed face-up at the top of a pole (45 cm length, 1.2 cm diameter), wrapped in tape for grip. A piece of cardboard was fitted over the top of the pole so that the mouse had to turn face down to descend the pole. Each assessment involved 2 days of habituation followed by 3 days of testing. An experimenter guided the mouse to complete the task during habituation. Testing was performed twice a day for three consecutive days and video was recorded. The recordings were later analyzed by an observer who was also blinded to the treatment conditions and scored for turn-around time (time taken to face downward) and descent time (time taken to come down the pole). However, descent-time was not used for data analysis because some impaired mice were unable to descend the pole normally and instead either slid down or used their tails to assist. Timing was begun with the first mouse movement, and any time exceeding 3 seconds during which the mouse was motionless was subtracted from the total time. Times were averaged for each mouse over the 3 trials.

### Brain harvest and tissue preparation

Mice were transcardially perfused with 200 ml of 0.9% cold saline and brains were removed. The brains were sagitally bisected. The right half was immersed in 4% paraformaldehyde in phosphate-buffered saline (PBS) for 48 hours, followed by immersion in 20% sucrose for 48 hours, and sectioned in a cryostat (40 μm thickness). The left half of the PFF bisected brains were stored at -80 °C until use for immunoblot preparation. At that time, a 1.0 mm coronal section was isolated from the frozen hemi-brain at the level of the substantia nigra and homogenized in PhosphoSure lysis buffer containing Hank’s phosphatase and protease inhibitor cocktail (ThermoScientific #78430). The tissue lysate was then sonicated on ice, and treated as described for the cell culture studies.

### Immunohistochemistry

Sections were rinsed in PBS, then incubated in blocking buffer (10% goat serum and 0.1% Triton X-100 in 0.1 M PBS) for 1 hour at room temperature. The sections were then incubated with designated primary antibodies overnight in blocking buffer. After washing, the sections were incubated with secondary antibodies that were either biotinylated or coupled to fluorescent tags. Biotinylated IgG secondary antibody was used at 5 μg / ml (Invitrogen, Carlsbad, CA) for 2-hour incubations, then processed using a Vector ABC kit (Vector laboratories). The horseradish peroxidase reaction was detected with diaminobenzidine and H_2_O_2_. Alexa Fluor secondary antibodies were used at 1:2000 dilutions for 2-hour incubations. After washing, the sections were mounted on slides with permount for DAB stained sections, or with ProLong™ Gold Antifade Mountant with or without DAPI (ThermoScientific, P36931, P36930) for fluorescent antibodies. Image analysis was done by an experimenter blinded to the experimental groups. Three to five photos were taken of the region of interest on each section. Identical camera settings were maintained for image capture within each experiment. In most studies the fluorescence intensity was measured as arbitrary units and normalized to either the number of cells or TH-positive cells in each field, as noted. In the studies using AAV, signals from the midbrain were normalized to signal from (non-transfected) cerebral cortex. For glutathione immunohistochemistry, 40 µm thick sections were incubated with 10 mM N-ethyl maleimide for 4 hours at 4 °C, washed, and incubated with mouse anti-glutathione-NEM (clone 8.1GSH, Millipore) as previously described and calibrated ^29^. After washing, the sections were incubated with Alexa Fluor 488-conjugated goat anti-mouse IgG (1:1000; Thermo Fisher Scientific) for 1 hour. Data from each animal were normalized to signal from cerebral cortex.

A stereological approach was used to assess tyrosine hydroxylase - positive neurons in the substantia nigra pars compacta. Three coronal sections were analyzed from each mouse, taken at the levels of -2.6, -3.2, and -3.6 mm anterior to Bregma, which nearly span the pars compacta ^30^. Tyrosine hydroxylase - positive cell bodies were counted in the substantia nigra pars compacta (as defined by Fu et al. ^30^) in each brain hemisection. The cell counts were made by an observer blinded to experimental conditions, and the counts for each mouse were expressed as the average number of cells per hemisection.

### Statistical analyses

For cell culture data, the “n” was defined as the number of experiments done using independent culture preparations, each with 3 - 4 internal replicates. For mouse studies, “n” denotes the number of animals in each group. When two groups were compared, data was analyzed by 2-sided student’s t test. For analyzing two or more groups, one-way ANOVA followed by the Bonferroni post hoc test for multiple comparisons. GraphPad Prism 8.0 software was used for all statistical analyses.

## RESULTS

We used cultured mouse neurons to investigate the relationships between α-synuclein aggregates, cell redox state, and c-Abl activation in cultured mouse neurons. Neurons were incubated with pre-formed human α-synuclein fibrils (PFFs) for 7 to 10 days to induce aggregation of endogenous α-synuclein ^20^. Aggregate formation was confirmed by detection of α-synuclein oligomers by western blotting and phospho-129 α-synuclein with immunostaining (Fig. 1A-D). No aggregation was observed in neurons prepared from α-synuclein deficient neurons (data not shown), consistent with prior reports ^20^. Aggregate formation was accompanied by oxidative stress, as shown by formation of the lipid peroxidation product 4-hydroxynonenal (Fig. 1C, D) and by conversion of dihydroethidium to oxidized ethidium species (Fig. 3B).

**Figure 1.**
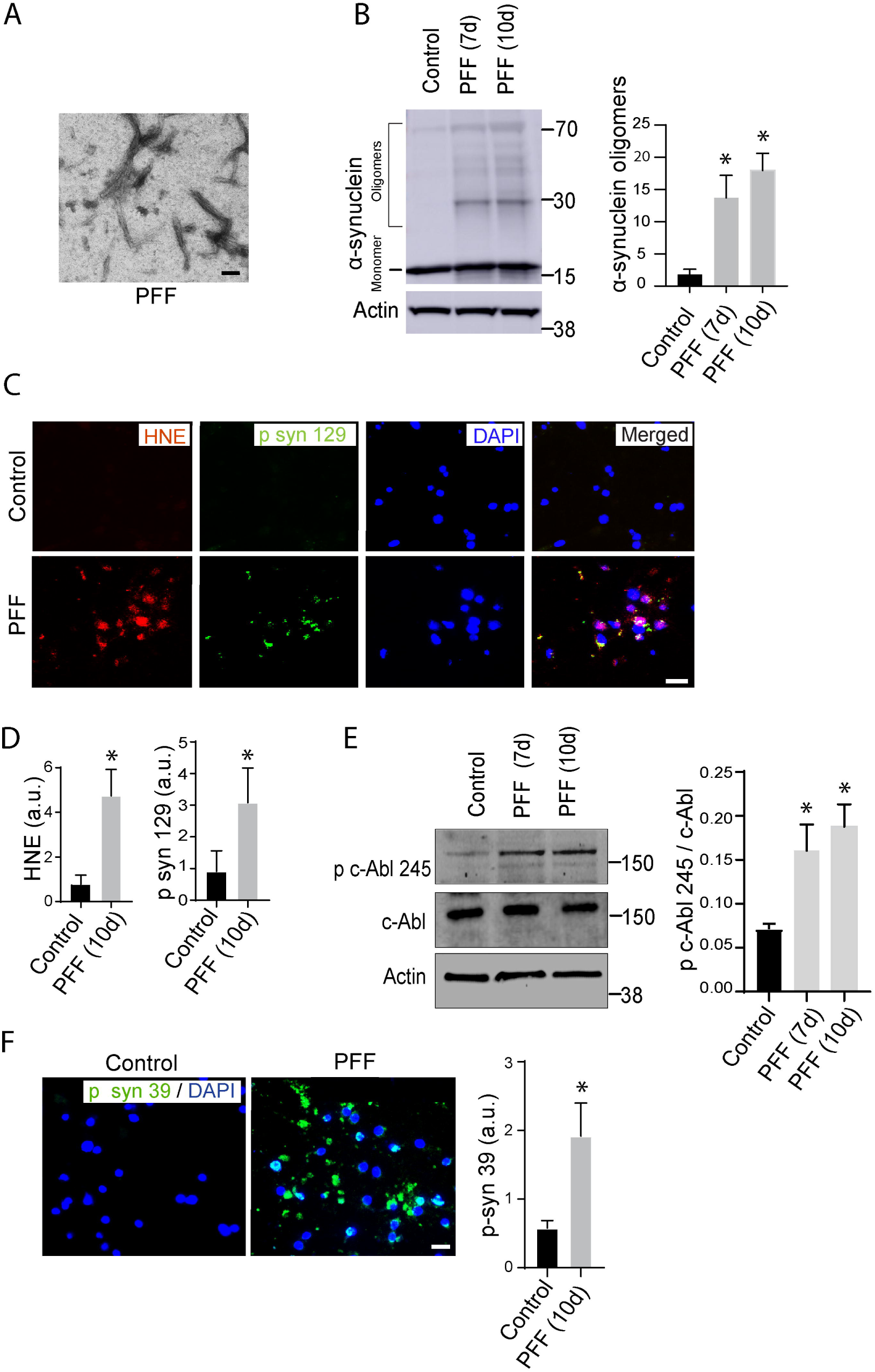
Pre-formed α-synuclein fibrils induce α-synuclein aggregation, oxidative stress, and c-Abl activation in cultured neurons. (**A)** Human α-synuclein pre-formed fibrils (PFFs) visualized by electron microscopy. Scale bar = 200 nm **(B)** Western blot showing formation of α-synuclein oligomers in primary cortical neurons treated with PFFs (5 µg / ml) for 7 or 10 days. Oligomers were quantified by summing the band densities over the molecular weight range indicated by bracket (20-75 Kda) and normalized to the corresponding actin loading control. n = 3, *p < 0.01 vs. control. **(C)** Immunostaining of primary cortical neurons treated with human PFFs for 10 days. α-synuclein aggregates are identified by p syn 129 (α-synuclein phosphorylated at serine129; green), oxidative stress is identified by the lipid oxidation marker HNE (4-hydroxynonenal; red), and cell nuclei are identified by DAPI (blue). Scale bar = 20 μm. The immunostaining is quantified in **(D)** with fluorescence expressed in arbitrary units and normalized to the number of cell nuclei in each field, n = 3, *p < 0.01. **(E)** Western blot shows c-Abl activation (c-Abl phosphorylation at tyrosine 245) in neurons treated with PFFs for 7 or 10 days. Quantification shows ratio of the phospho c-Abl band to total c-Abl. n = 3, *p < 0.01 vs. control. **(F)** Immunostaining for p-syn 39 (α-synuclein phosphorylated at tyrosine 39; green) in neurons treated with PFFs for 10 days. DAPI (blue) labels cell nuclei. Scale bar = 20 μm. Graph shows fluorescence expressed in arbitrary units and normalized to the number of cell nuclei in each field. n = 3, *p < 0.01 vs control.

**Figure 2.**
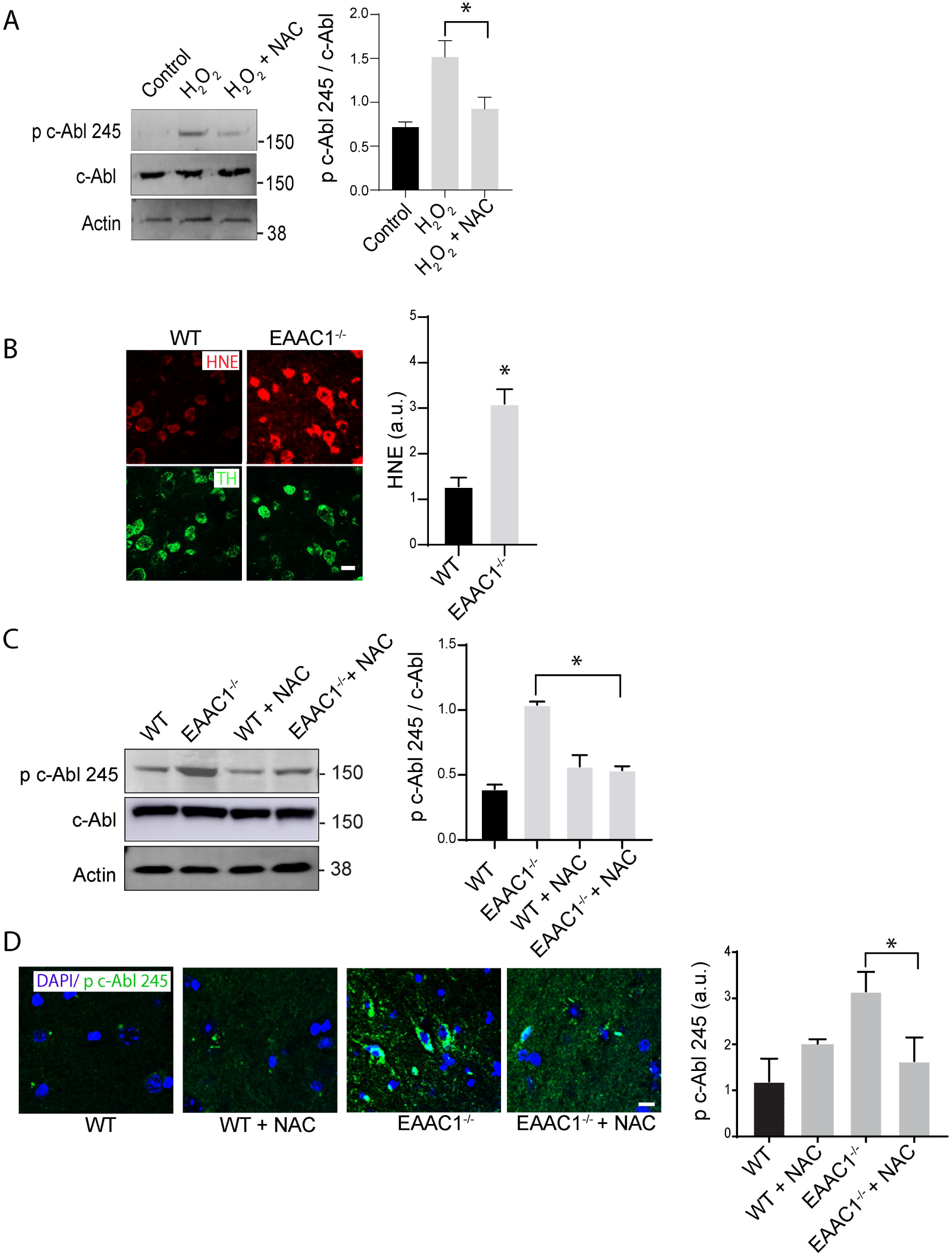
Oxidative stress alone is sufficient to activate c-Abl in culture and *in vivo.* **(A)** Western blot shows activation (phosphorylation) of c-Abl in neurons treated with 100 µM hydrogen peroxide for 30 minutes, and reduced activation in the additional presence of 500 µM N-acetyl cystine (NAC). Quantification shows ratio of the phospho c-Abl band to total c-Abl. n = 3, *p < 0.01 **(B)** Oxidative stress in TH neurons (green) as detected by 4-hydroxynonenal (HNE, red) in EAAC1^-/-^ mouse midbrain. Scale bar = 20 μm. Quantified HNE fluorescence present in TH cells. n = 3, * p < 0.01 WT vs EAAC1^-/-^. **(C)** Western blot showing activation (phosphorylation) of c-Abl in midbrain of EAAC1^-/-^ mice and suppression of this activation in EAAC1^-/-^ mice treated with NAC for 10 days. Quantification shows ratios of the phospho c-Abl band to total c-Abl, n = 3; *p < 0.01. (**D)** Immunostaining showing increased phospho c-Abl (green) in midbrain of EAAC1^-/-^ mice and reversal of this increase in EAAC1^-/-^ mice treated with NAC for 10 days. Cell nuclei are labeled with DAPI. Scale bar = 10 μm. Quantified phospho c-Abl, n = 3, *p < 0.01.

**Figure 3.**
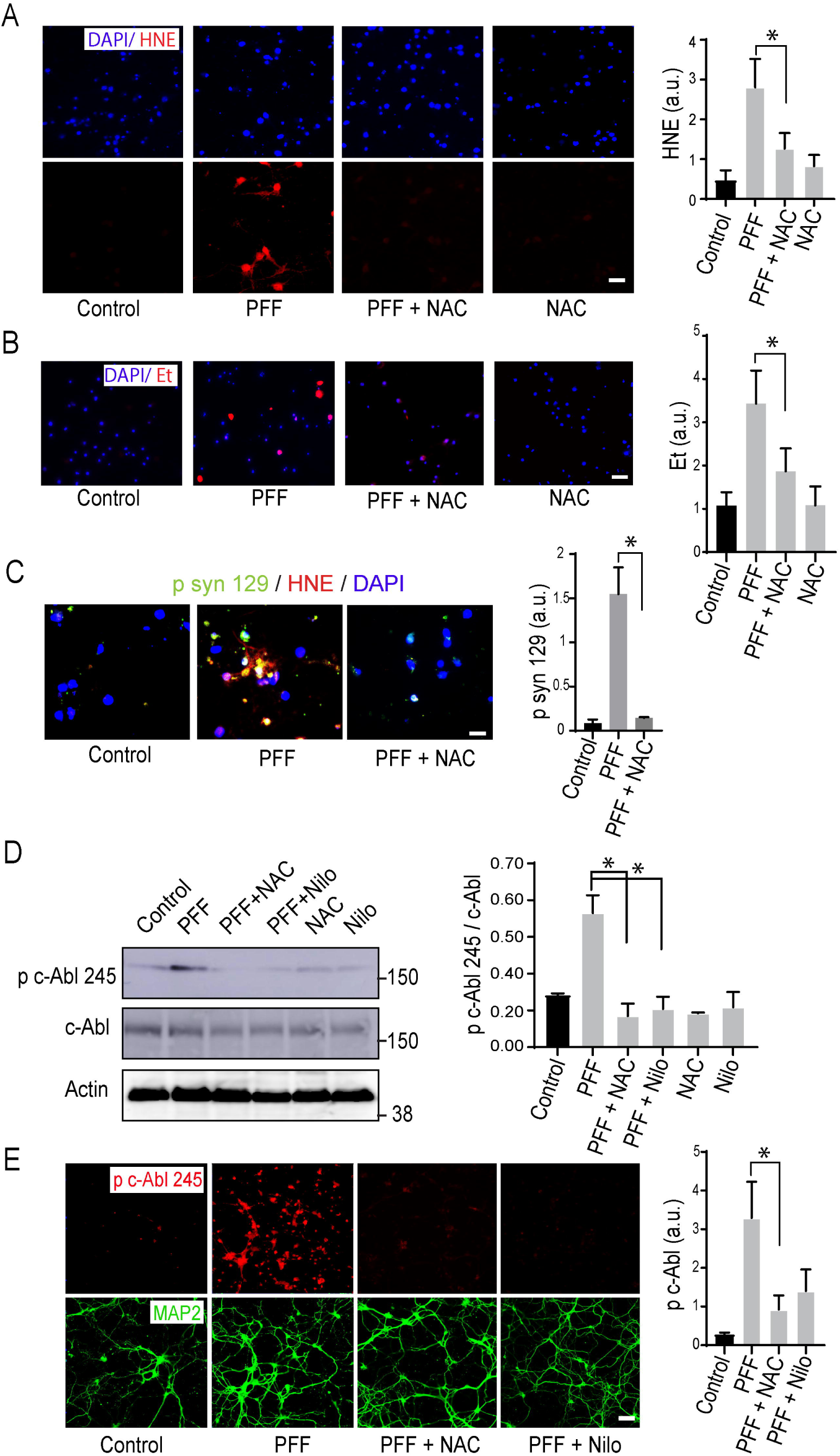
N-acetyl cysteine suppresses oxidant stress and c-Abl activation in cultured neurons. (**A**) Oxidant stress detected by formation of 4-hydroxynonenal (HNE, red) in neurons incubated with PFFs for 10 days with or without 500 µM N-acetyl cysteine (NAC). Cell nuclei are labeled blue with DAPI. Scale bar = 20 μm. Quantified HNE signal is expressed in arbitrary units and normalized to the number of cell nuclei in each field. n = 3, * p < 0.01 (**B**) Oxidant stress detected by formation of fluorescent ethidium species (Et, red) in neurons incubated with PFFs for 10 days with or without 500 µM NAC. Cell nuclei are stained blue with DAPI. Scale bar = 20 μm. Quantified Et signal is expressed in arbitrary units and normalized to the number of cell nuclei in each field. n = 3, * p < 0.01. (**C**) α-synuclein aggregation as detected by p syn 129 immunostaining (green) in neurons incubated with PFFs for 10 days with or without addition of 500 µM N-acetyl cysteine (NAC). Oxidant stress detected by formation 4-hydroxynonenal (HNE, red) and cell nuclei are labeled with DAPI (blue). Scale bar = 10 μm. Quantified p syn 129 signal is expressed in arbitrary units and normalized to the number of cell nuclei in each field. n = 3, * p < 0.01 (**D**) Western blot shows activation (phosphorylation) of c-Abl in neurons treated 10 days with PFFs, and reduced activation in the additional presence of 500 µM NAC or 10 µM nilotinib. Quantification shows ratios of the phospho c-Abl band to total c-Abl, n = 3, *p < 0.01 (**E**) Immunostaining shows c-Abl activation (c-Abl phosphorylation at tyrosine 245; red) in neurons treated 10 days with PFFs, and reduced activation in the additional presence of 500 µM NAC or 10 µM nilotinib. MAP2 (green) identifies neuronal cytoplasm. Scale bar = 20 μm. Graph shows p c-Abl 245 fluorescence expressed in arbitrary units and normalized to the neuronal (MAP2) area in each field, n = 3, *p < 0.01.

We assessed c-Abl activation in the PFF-exposed cultures by western blotting with antibodies to total c-Abl and activated c-Abl (phosphorylated at tyrosine 245) ^14^. Neurons incubated with PFFs showed a large increase in activated c-Abl, with no change in total c-Abl (Fig. 1E). c-Abl activation in the PFF-exposed cells was further evidenced by a large increase in phospho-39 α-synuclein (Fig. 1F), which is produced by activated c-Abl ^17^. Together, these results show that PFF-induced α-synuclein aggregates produce both oxidative stress and c-Abl activation in cultured neurons.

### Oxidative stress alone is sufficient to activate c-Abl in culture and *in vivo*

To determine whether oxidative stress is sufficient to cause c-Abl activation in neurons, we exposed cultures to hydrogen peroxide. Western blots prepared from these cultures showed a robust increase in c-Abl activation (Fig. 2A). The H_2_O_2_ - induced c-Abl activation was suppressed by N-acetyl-cysteine (NAC), a thiol agent previously shown to support neuronal glutathione synthesis ^28, 31^ and to have salutary effects in animal models of PD ^28, 32, 33^. We next evaluated the effect of redox stress on c-Abl activation *in vivo*, using EAAC1^-/-^ mice. EAAC1 (also termed EAAT3 and SLC1A1) is a neuronal cysteine transporter, and mice deficient in EAAC1^-/-^ mice have reduced glutathione levels and chronic oxidative stress in neurons ^25, 28^ (Fig. 2B). Western blot and immunofluorescence measures of phospho c-Abl in EAAC1^-/-^ mouse cerebral cortex midbrain showed increased c-Abl activation, and additionally showed this to be prevented by administration of NAC (3mg / ml) in drinking water (Fig. 2C, D), thus demonstrating that chronic oxidative stress is sufficient to drive neuronal c-Abl activation.

### N-acetyl cysteine suppresses oxidant stress and c-Abl activation in cultured neurons

If the c-Abl activation induced by α-synuclein aggregates is mediated by oxidative stress, then suppression of oxidative stress should attenuate c-Abl activation in the PFF-treated neurons. This was tested by incubating the PFF - treated neuronal cultures with NAC. NAC reduced formation of both HNE and oxidized ethidium species (Fig 3A, B), and also reduced formation of α-synuclein aggregates (Fig. 3C). NAC also suppressed c-Abl activation, as demonstrated both by immunostaining and western blots (Fig. 3D, E). The effect of NAC was comparable to that achieved by the c-Abl inhibitor nilotinib, which is a targeted inhibitor of c-Abl ^34^ and included here as a positive control.

### PFFs injections induce oxidative stress and c-Abl activation *in vivo*

We next evaluated effects of PFF-induced oxidative stress and c-Abl activation in mouse brain. Mice aged 9 - 10 months received bilateral injections of PFFs (or saline control) into the substantia nigra ^35^. NAC was added to the drinking water of half of the PFF-injected mice beginning 2 weeks after the injections and continued until brain harvest 6 months later. Assessment of the substantia nigra in these brains showed that PFF injections reduced neuronal glutathione content (indicative of redox stress) and that NAC attenuated this effect (Fig. 4). PFF injections also produced α-synuclein aggregation and lipid peroxidation in substantia nigra neurons (Fig. 4). The aggregates and lipid peroxidation were prominent in dopaminergic (tyrosine hydroxylase (TH) - positive) neurons, but not restricted to these cells. Mice treated with NAC exhibited a substantial reduction in lipid peroxidation (Fig. 4B, C), indicating attenuated redox stress. These mice also showed suppression of c-Abl activation, as assessed by both phospho c-Abl formation and phospho-synuclein 39 formation (Fig. 4D, E), thus indicating redox stress as the primary factor driving c-Abl activation. Surprisingly, there was also a substantial reduction in α-synuclein aggregate formation in the mice treated with NAC (Fig. 4B, C). Evaluation of TH - positive cells in the substantia nigra pars compacta showed a reduction in mice subjected to PFF injections, and this was accompanied by reduced performance on the rota-rod test and increased turn-around time on the pole test (Fig. 5). These impairments were also attenuated in the mice treated with NAC.

**Figure 4.**
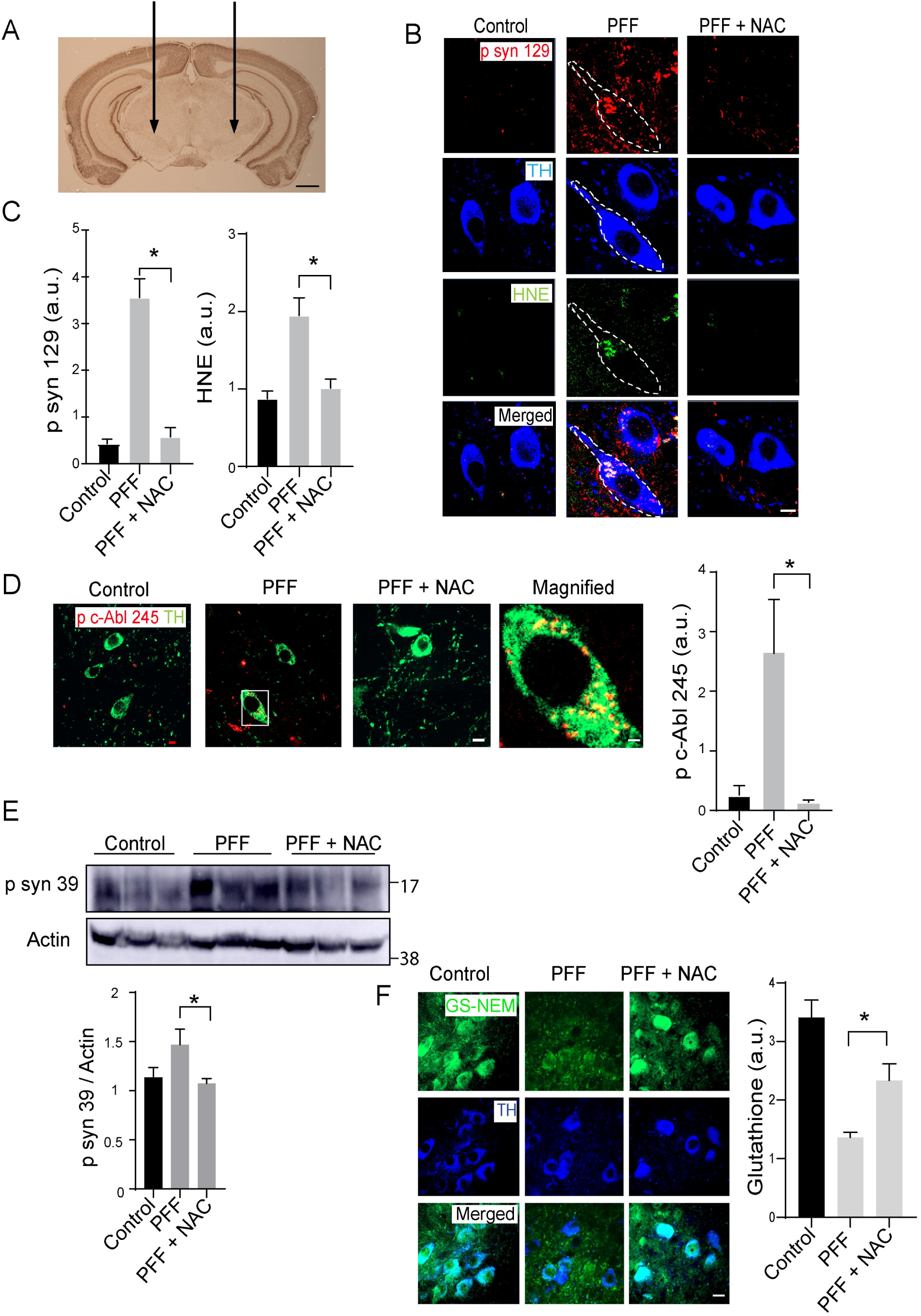
PFFs induce c-Abl activation *in vivo.* **(A)** Coronal mouse section indicating sites of stereotactic PFF injections (substantia nigra). **(B)** Immunostaining at the injection site identifying dopaminergic cell bodies (TH; blue), α-synuclein aggregates (p syn 129; red), and lipid peroxidation (HNE; green). Brains were harvested 6 months after PFF injections. Where indicate, the mice also received 3 mg/ml NAC in drinking water beginning 2 weeks post-injection. Scale bar = 8 μm. **(C)** Quantification of p syn 129 and HNE fluorescence in the TH positive dopaminergic neurons, expressed as arbitrary units. n = 3 - 4, *p < 0.01. **(D)** Immunostaining for activated c-Abl (p c-Abl 245; red) in TH - positive (green) cells. Scale bar = 5 μm. White rectangle denotes field shown in magnified view. Quantified data shows phospho c-Abl signal fluorescence in TH cells. n = 3-4; *p < 0.01 (**E)** Western blot shows phosphorylation of synuclein at tyrosine 39 in midbrain lysates obtained from control, PFF-injected, and PFF-injected mice treated with NAC. Quantification shows ratio of the p-syn 39 band to actin n = 3, *p < 0.05 **(F)** Immunostaining for glutathione (green) in TH - positive (blue) cells. Scale bar = 10 μm. Quantified data are from n = 3 - 4; *p < 0.05.

**Figure 5.**
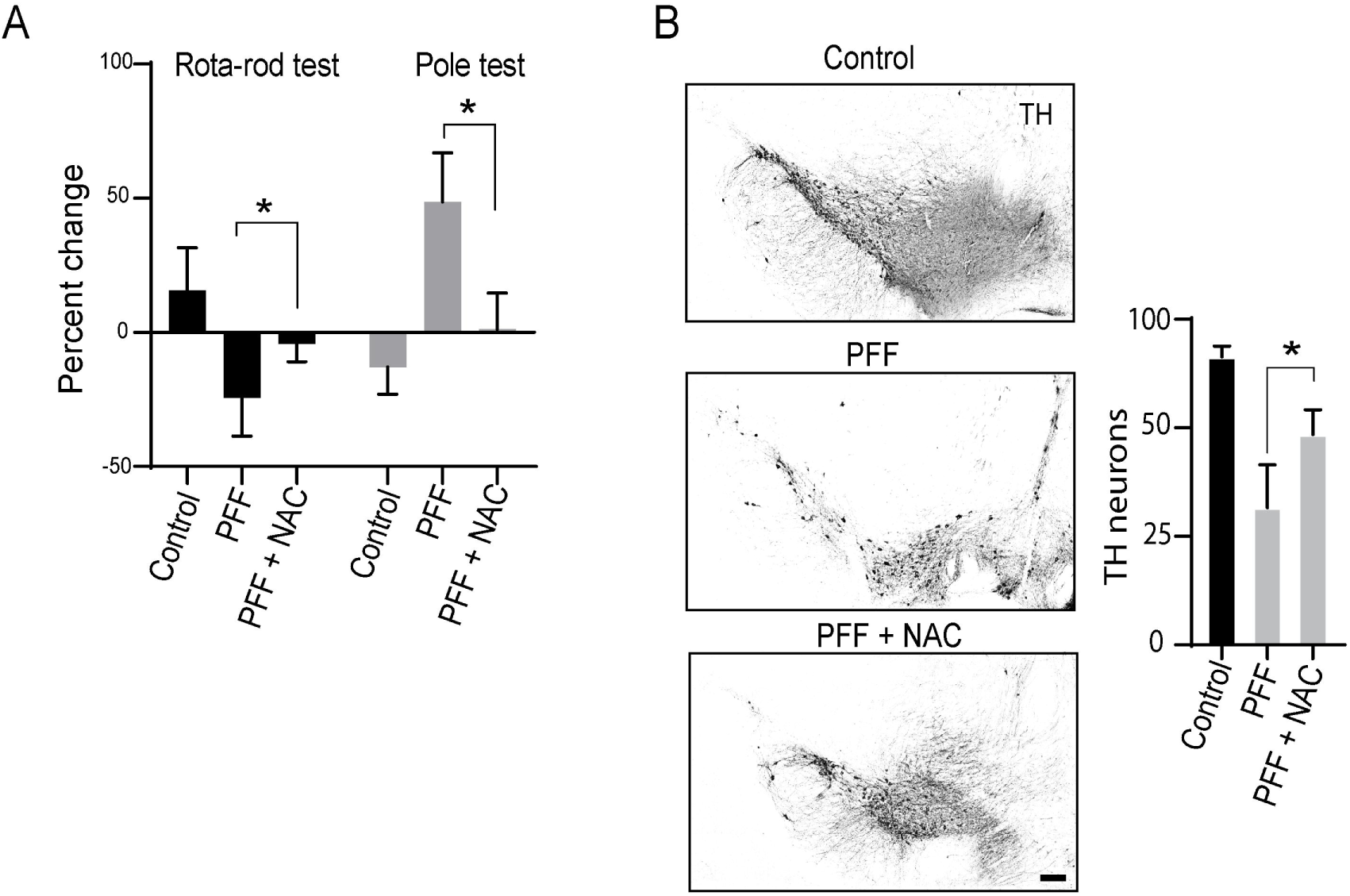
NAC improves neuron survival and motor function in PFF-injected mice. **(A)** Performance on rota-rod and pole test 4 months after injections. n = 3 - 4; *p < 0.01. **(B)** TH-positive cells in substantia nigra of control, PFF-injected and PFF-injected mice treated with NAC. Scale bar = 50 μm. Number of TH neurons in substantia nigra pars compacta from each hemisphere were quantified. n = 3 - 4; *p < 0.01

### AAV-mediated α-synuclein overexpression induces oxidative stress and c-Abl activation

As a second mouse model of α-synuclein aggregate formation, we injected adeno-associated virus expressing human α-synuclein into the substantia nigra ^36^. Controls were injected with AAV expressing mCherry (Fig 6A). NAC was added to the drinking water of half of the mice beginning 2 weeks after the AAV injections and continued until brain harvest 6 months later. Assessment of the substantia nigra in these brains showed that the AAV α-synuclein injections produced α-synuclein aggregates and oxidative stress, along with c-Abl activation (Fig. 6B, C). Co-labeling for TH revealed phospho (activated) c-Abl in dopaminergic neurons (Fig. 6D). Dopaminergic cell death could not be assessed in the substantia nigra of these mice because of insufficient material for stereological assessment. However, immunostaining for dopaminergic projections into the striatum showed a decrease in TH - positive innervation in brains injected with AAV-α-synuclein, and an attenuation of this decrease by NAC treatment (Fig. 6E). As with the PFF injections, mice injected with AAV-α-synuclein also exhibited reduced motor dexterity as assessed by the rota-rod test and pole test, which were also attenuated by NAC treatment.

**Figure 6.**
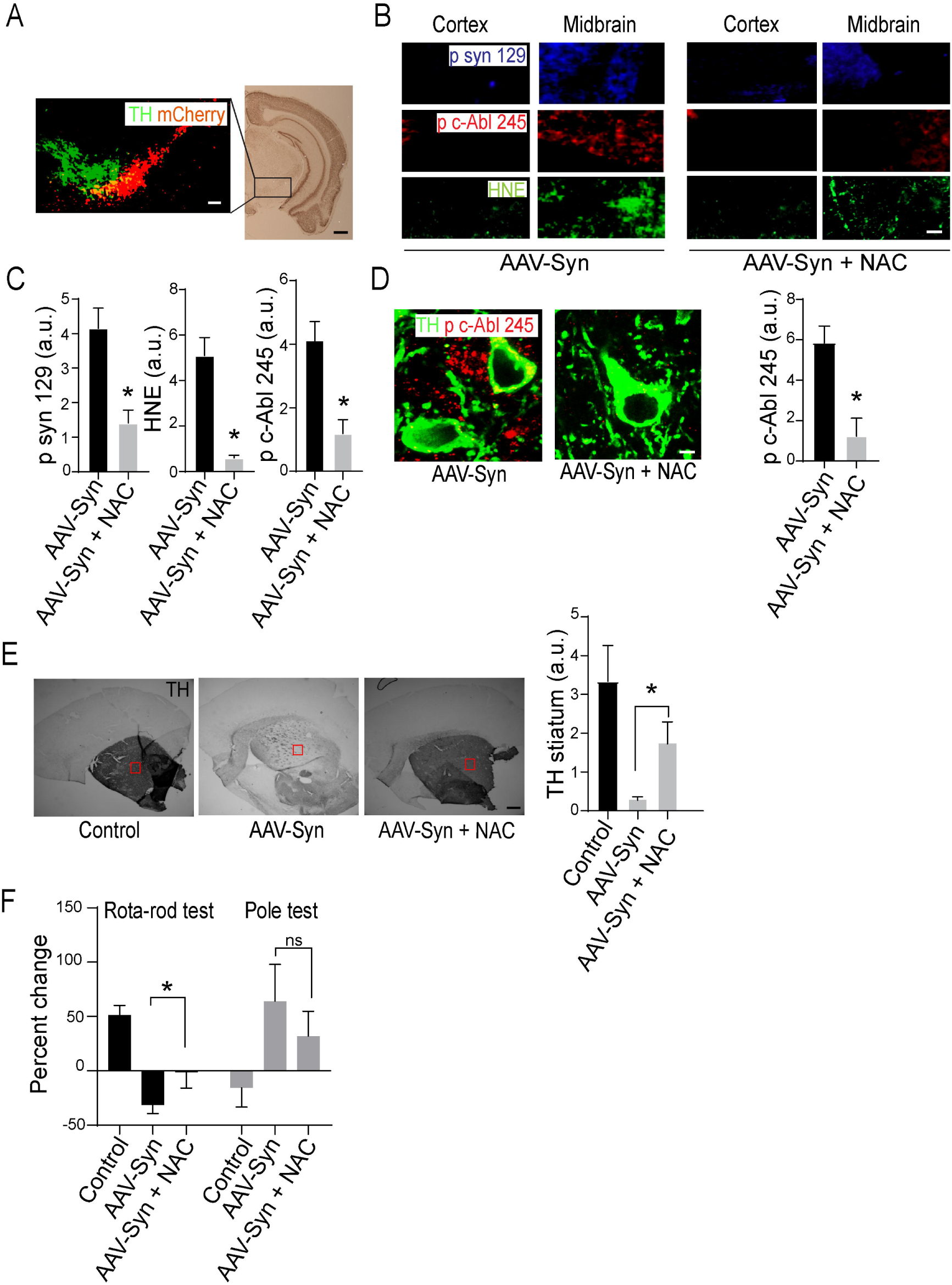
NAC improves outcome after AAV-mediated α-synuclein overexpression. **(A)** Coronal hemisection showing site of AAV injection and virus expression (red; mCherry). Immunostaining for TH (green) identifies dopaminergic neurons. Scale bar = 250 μm. **(B** Images taken from midbrain and cerebral cortex showing HNE (green), phospho c-Abl (red) and phospho syn 129 (blue) in mice 6 months after injection with α-synuclein AAV, with or without subsequent oral NAC treatment. Scale bar = 40 μm **(C)** Quantification of midbrain HNE, c-Abl and phospho syn 129 signal, expressed relative to signal in cortex, n = 3 - 4; *p < 0.01 **(D)** p c-Abl 245 (red) in TH positive cells (green) from substantia nigra of mice injected with α-synuclein AAV with or without subsequent NAC treatment. Scale bar = 3 μm Quantified results are from n = 3 - 4; *p < 0.01 **(E)** DAB immunostaining of TH in striatum of mice injected with α-synuclein AAV-syn with and without subsequent NAC treatment. Scale bar = 250 μm. Quantified results are from n = 3 - 4; *p < 0.01 **(F)** Performance on rota-rod test and pole test 5 months after AAV injections. n = 4 - 6; *p < 0.01.

## DISCUSSION

Interactions between cell redox state and α-synuclein may underlie convergent mechanisms of disease progression in PD. Our findings show that α-synuclein aggregates induce c-Abl activation by a process involving redox stress. Given that both oxidative stress ^3, 5, 6^ and c-Abl activation ^14, 15, 17^ promote α-synuclein aggregate formation, these results identify a potentially feed-forward process (Fig. 7). Consistent with this idea, our findings also show that NAC suppresses the oxidant stress caused by α-synuclein aggregates, and that this likewise suppresses both c-Abl activation and aggregate formation itself.

**Figure 7.**
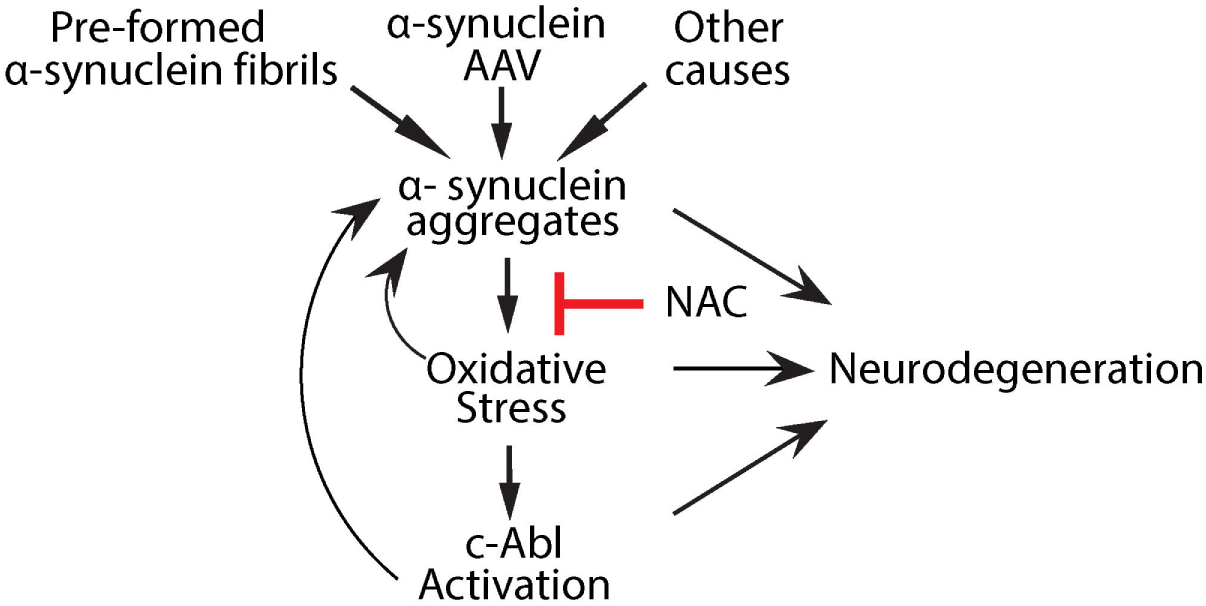
Proposed Interactions between α-synuclein aggregates, oxidative stress, and c-Abl. α-synuclein aggregates produce oxidative stress, which drives c-Abl activation. Oxidative stress and c-Abl activation both contribute to α-synuclein aggregation, in a potentially feed-forward process. NAC suppresses the oxidative stress induced by α-synuclein aggregates and thereby attenuates c-Abl activation and further α-synuclein aggregation.

Large, histochemically identifiable aggregates of α-synuclein (Lewy bodies) are a diagnostic feature of PD. These large aggregates are unlikely to be directly cytotoxic, but they are strongly associated with cytotoxicity that is putatively mediated by smaller, oligomeric aggregates or by products cleaved from aggregates ^37^, or by sequestration of soluble α-synuclein ^26^. Aggregate formation can be induced experimentally by exposure to pre-formed human α-synuclein fibrils (PFFs) through a putative prion-like mechanism ^20, 38^. Here we used PFFs to induce formation of α-synuclein aggregates in neuronal cultures and *in vivo*. To complement the studies performed with PFFs, we also used an AAV-mediated α-synuclein overexpression approach, which likewise induces aggregate formation ^39^. Both studies employed relatively old mice, age 9 - 15 months, which are more prone to develop α-synuclein aggregates and better mimic human PD. Aggregates induced by both approaches produced redox stress and c-Abl activation. They also produced dopaminergic neurodegeneration and associated motor deficits. The severity of the motor deficits was modest, consistent with prior studies indicating relative preservation of motor function in mice with incomplete or unilateral injury to substantia nigra ^40^.

c-Abl activation promotes the formation or stability of α-synuclein aggregates, and may thereby contribute to PD pathogenesis ^13^. However, the trigger for c-Abl activation in PD has been uncertain. Cell culture studies cells indicate that oxidative stress can activate c-Abl by direct or indirect pathways ^11-13^. Our findings confirm that oxidant stress is sufficient to induce c-Abl, both in neuronal cell cultures with acute exposure of H_2_O_2_, and *in vivo* using the EAAC1^-/-^ mouse of chronic neuronal redox stress. The neuronal glutathione deficit and other phenotypic features of EAAC1^-/-^ mice can be negated by oral NAC administration ^28^, and the present results show that NAC treatment also blocks c-Abl activation in these mice.

The mechanism by which α-synuclein aggregates lead to redox stress was not evaluated in this study. Prior work suggests that this may occur through the interaction of aggregates or aggregate cleavage products with mitochondria ^41, 42^ or transition state metals in neuronal cytosol ^7^. It is also possible that aggregates induce activation of NADPH oxidase, which produces superoxide and has been identified as a potential pathogenic factor in PD ^43^.

Although NAC suppressed c-Abl activation both in cell culture and *in vivo*, the mode of NAC action may differ in the two settings. NAC administration *in vivo* supports synthesis of neuronal glutathione, which is used as a cofactor for enzyme-mediated protein repair and oxidant scavenging. Most glutathione is recycled, and consequently low CSF concentrations of NAC are sufficient to maintain neuronal glutathione levels ^31^. As demonstrated in the present study, 3 mg / ml of NAC in water was sufficient to maintain normal glutathione levels in neurons containing α-synuclein aggregates. Neurons in culture have ready access to cysteine and cystine in the culture medium, and in this setting the anti-oxidant effect of NAC is independent of glutathione synthesis and attributable to direct oxidant scavenging by its reactive cysteine thiol, which requires a higher NAC concentration ^44^. NAC may also have additional actions on neurons mediated by its effects on cell redox state or ferroptosis ^45^.

NAC also reduced formation of α-synuclein aggregates both in the cell culture and the *in vivo* models of PD. While the reduced aggregate formation is relevant to PD therapeutics, this also makes it difficult to ascertain whether the associated reduction in oxidant stress and dopaminergic neurodegeneration were direct effects of NAC, or indirect effects mediated by the reduced aggregate formation. However, given the feed-forward relationships between these events (Fig. 7), a distinction between these two alternatives may not be meaningful.

Glutathione levels are decreased selectively in the substantia nigra pars compacta (SNc) of patients with PD and presumed pre-symptomatic PD ^46, 47^, and studies using animal models suggest that this reduction contributes to disease progression ^28, 48, 49^. In accordance with this idea, NAC has been shown to have beneficial effects in both toxin and genetic models of PD ^28, 32, 33^. NAC is inexpensive, FDA-approved, orally absorbed, and crosses the blood-brain barrier ^28, 31^. Thiol repletion using NAC or other agents has been repeatedly cited as a promising approach for slowing PD progression ^31, 46, 50^, and results of the present study provide additional mechanistic support for this approach.

## ACKNOWLEDGEMENTS

We thank Robert Edwards, University of California San Francisco, for the α, β, γ synuclein triple knockout mice. This work was supported by the U.S. National Institutes of Health (NS105774, R.A.S.; DK108798, F. M.) and Dept. of Veterans Affairs (BX003249, R.A.S.).

## AUTHOR CONTRIBUTIONS

S.G and R.A.S contributed to the conception and design of the study; S.G., S.J.W., R. F., N.J.M.B., J.P., J.S., L.W., J.W. and F.P.M. contributed to the acquisition and analysis of data; S.G. and R.A.S. contributed to drafting the text and preparing the figures.

